# Elevated polygenic burden for autism is associated with differential DNA methylation at birth

**DOI:** 10.1101/225193

**Authors:** Eilis Hannon, Diana Schendel, Christine Ladd-Acosta, Jakob Grove, iPSYCH-Broad ASD Group, Christine Søholm Hansen, Shan V. Andrews, David Michael Hougaard, Michaeline Bresnahan, Ole Mors, Mads Vilhelm Hollegaard, Marie Bækvad-Hansen, Mady Hornig, Preben Bo Mortensen, Anders D. Børglum, Thomas Werge, Marianne Giørtz Pedersen, Merete Nordentoft, Joseph Buxbaum, M Daniele Fallin, Jonas Bybjerg-Grauholm, Abraham Reichenberg, Jonathan Mill

**Affiliations:** University of Exeter Medical School, University of Exeter, UK; Department of Public Health, Aarhus University, Aarhus, Denmark; Department of Epidemiology and Johns Hopkins Bloomberg School of Public Health, Baltimore, MD, USA; Wendy Klag Center for Autism and Developmental Disabilities, Johns Hopkins Bloomberg School of Public Health, Baltimore, MD, USA; Department of Biomedicine, Aarhus University, Aarhus, Denmark; iPSYCH, The Lundbeck Foundation Initiative for Integrative Psychiatric Research, Denmark; Centre for Integrative Sequencing, iSEQ, Aarhus University, Aarhus, Denmark; Bioinformatics Research Centre, Aarhus University, Aarhus, Denmark; Center for Neonatal Screening, Department for Congenital Disorders, Statens Serum Institut, Copenhagen, Denmark; Institute of Biological Psychiatry, MHC Sct. Hans, Mental Health Services Copenhagen, Roskilde, Denmark; Center for Infection and Immunity, Columbia University Mailman School of Public Health, New York, USA; Department of Epidemiology, Columbia University Mailman School of Public Health, New York, USA; Psychosis Research Unit, Aarhus University Hospital, Risskov, Denmark; Department of Clinical Medicine, Aarhus University; Aarhus University Hospital, Risskov; National Centre for Register-Based Research, Aarhus University, Aarhus, Denmark; Centre for Integrated Register-based Research, Aarhus University, Aarhus, Denmark; Department of Clinical Medicine, University of Copenhagen, Copenhagen, Denmark; Mental Health Services in the Capital Region of Denmark, Mental Health Center Copenhagen, University of Copenhagen, Copenhagen, Denmark; Department of Psychiatry, Mount Sinai School of Medicine, USA; Department of Psychiatry, Columbia University, New York, USA; Department of Mental Health, Johns Hopkins Bloomberg School of Public Health, Baltimore, MD USA

**Keywords:** Autism, DNA methylation, genetics, neonatal, genome-wide association study (GWAS), epigenome-wide association study (EWAS), birth, DNA methylation quantitative trait loci (mQTL)

## Abstract

**Background:** Autism spectrum disorder (ASD) is a severe neurodevelopmental disorder characterized by deficits in social communication and restricted, repetitive behaviors, interests, or activities. The etiology of ASD involves both inherited and environmental risk factors, with epigenetic processes hypothesized as one mechanism by which both genetic and non-genetic variation influence gene regulation and pathogenesis.

**Methods:** We quantified neonatal methylomic variation in 1,263 infants - of whom ~50% went on to subsequently develop ASD – using DNA isolated from a unique collection of archived blood spots taken shortly after birth. We used matched genetic data from the same individuals to examine the molecular consequences of ASD genetic risk variants, identifying methylomic variation associated with elevated polygenic burden for ASD. In addition, we performed DNA methylation quantitative trait loci (mQTL) mapping to prioritize target genes from ASD GWAS findings.

**Results:** Although we did not identify specific loci showing consistent changes in neonatal DNA methylation associated with later ASD, we found a significant association between increased polygenic burden for autism and methylomic variation at two CpG sites located proximal to a robust GWAS signal for ASD on chromosome 8.

**Conclusions:** This study is the largest analysis of DNA methylation in ASD yet undertaken and the first to integrate both genetic and epigenetic variation at birth in ASD. We demonstrate the utility of using a polygenic risk score to identify molecular variation associated with disease, and of using mQTL to refine the functional and regulatory variation associated with ASD risk variants.

## BACKGROUND

Autism spectrum disorder (ASD) defines a group of complex neurodevelopmental disorders marked by deficits in social communication and restricted, repetitive behaviors, interests, or activities[1].

ASD affects ~1–2% of the population, and confers severe lifelong disability[2–4]. Quantitative genetic studies indicate that ASD is highly heritable[5, 6], although population-based epidemiologic studies of environmental risks and ASD liability modelling using family designs also indicate environmental factors as important[7]. Genetic studies have shown that autism risk is strongly associated with both rare inherited and *de novo* DNA sequence variants[8–11]. In contrast, the identification of common genetic variants associated with ASD using genome-wide association studies (GWAS) has proven harder than for other complex neuropsychiatric traits such as schizophrenia[12], at least in part due to a lack of large sample datasets. Recent collaboration between the Psychiatrics Genomics Consortium autism workgroup (PGC-AUT) and the Lundbeck Foundation Initiative for Integrative Psychiatric Research (iPSYCH) has greatly expanded the number of ASD cases with GWAS data, identifying three genome-wide significant associations for ASD and evidence for a substantial polygenic component in signals falling below the stringent genome-wide significance threshold (Grove et al, https://www.biorxiv.org/content/early/2017/11/25/224774). None of the three ASD-associated loci are predicted to result in coding changes or altered protein structure; instead they are hypothesized to influence gene regulation. Previous studies of other neurodevelopmental disorders have reported an enrichment of disease-associated variation in regulatory domains, including enhancers and regions of open chromatin[13].

Epigenetic variation induced by non-genetic exposures has been hypothesized to be one mechanism by which environmental factors can affect risk for ASD[14, 15]. Recent studies have provided initial evidence for autism-associated epigenetic variation in both brain and peripheral tissues[16–21], although these analyses have been undertaken on relatively small numbers of samples with limited statistical power. Existing analyses have assessed epigenetic variation in samples collected after a diagnosis of ASD has been assigned and are potentially confounded by factors such as smoking[22–24], medication [25, 26], other environmental toxins[27] and reverse causation[28]. Furthermore, they have not investigated the role of genetic variation in mediating associations between epigenetic variation and ASD. The integration of genetic and epigenetic data will facilitate a better understanding of the molecular mechanisms involved in autism, especially given the high heritability of ASD and recent data showing how the epigenome can be directly influenced by genetic variation[29–32]. For example, we have previously demonstrated the potential for using polygenic risk scores (PRS) - defined as the sum of trait-associated alleles across many genetic loci, weighted by GWAS effect sizes - as disease biomarkers with utility for exploring the molecular genomic mechanisms involved in disease pathogenesis [33]. Of note, PRS-associated epigenetic variation is potentially less affected by factors associated with the disease itself, which can confound case–control analyses.

In this study, we quantified DNA methylation for ~1,316 individuals (comprising equal numbers of ASD cases and matched controls, 50% male/female) isolated from neonatal blood spots collected proximal to birth (mean = 6.08 days; sd = 3.24 days; **Supplementary Figure 1**). Known epigenetic signatures for gestational and chronological age[34, 35], and exposure to maternal smoking during pregnancy[23], were used to confirm the robust nature of genome-wide DNA methylation data generated from neonatal blood spots. Matched genome-wide single nucleotide polymorphism (SNP) genotyping data from the same individuals enabled us to undertake an integrated genetic-epigenetic analysis of ASD, exploring the extent to which neonatal methylomic variation at birth is associated with elevated polygenic burden for ASD. Finally, we generated an extensive database of DNA methylation quantitative trait loci (mQTL) in neonatal blood samples, which were used to characterize the molecular consequences of genetic variants associated with ASD.

## METHODS

### Overview of the MINERvA cohort

Denmark has a comprehensive neonatal screening program which is used to test for innate errors of metabolism, hypothyroidism and other treatable disorders. Neonatal blood is collected on standard Guthrie cards and residual material is stored within the Danish Neonatal Screening Biobank. The reason for storing the samples in prioritized order is: (1) diagnosis and treatment of congenital disorders, (2) diagnostic use later in infancy after informed consent, (3) legal use after court order, (4) research projects pending approval by the Scientific Ethical Committee System in Denmark, The Danish Data Protection Agency and the NBS-Biobank Steering Committee. Thus, research is possible assuming sufficient material remains for the proceeding priorities[36]. Cases and controls were selected from the iPSYCH case-control sample, which has been recently described[37]. Briefly, the iPSYCH study population comprises all singletons born in Denmark between May 1st 1981 and December 31st 2005, who are still alive and residing in Denmark at their first birthday and with a known mother. iPSYCH ASD cases comprise all children in the study population with an ASD diagnosis reported before December 31st 2012. iPSYCH controls comprise 30,000 persons randomly selected from the study population (about 2% of the total study population).

The MINERvA study profiled a subsample of 1,316 iPSYCH samples, including an equal number of ASD cases and controls that were selected using the following criteria. Cases were born between 1998 and 2002, with both parents born in Denmark themselves. We selected a 1:1 male to female ratio (i.e. by ‘oversampling’ ASD females). Cases and controls were excluded if they had a reported diagnosis (before December 31st 2012) of select known genetic disorders: Down syndrome, Fragile X, Angelman, Prader Willi, Zellweger, William, tuberous sclerosis, Rett, Tourette, neurofibromatosis, Duchennes, Cornelia de Lange, DiGeorge, Smith-Lemli-Opitz, Klinfelter. In addition, controls were excluded if they had died or emigrated from Denmark before December 31st 2012, or had any reported psychiatric diagnosis. Eligible controls were individually matched to cases on sex, month of birth (month before, same month, or month after case month) and year of birth. Among the controls fulfilling these criteria, additional matching criteria were applied as closely as possible with regard to gestational age (in weeks) and the same urbanicity level of maternal residence at time of birth as cases. All perinatal data used for case-control matching, plus additional information on birth weight and maternal smoking were obtained from the Danish Medical Birth Register or the Central Person Register. Detailed maternal smoking data was used to generate a binary variable indicating whether the mother smoked during pregnancy or not. All diagnoses used for ASD case identification and case/control exclusions were obtained from the Danish Psychiatric Central Research Register (DPCRR) and Danish National Patient Register (DNPR). In Denmark, children and adolescents suspected of ASD or other mental or behavioral disorders are referred by general practitioners or school psychologists to a child and adolescent psychiatric department for a multidisciplinary evaluation, and their conditions are diagnosed by a child and adolescent psychiatrist. Registry reporting is done only by psychiatrists following mandatory training in the use of the World Health Organization International Classification of Diseases (ICD)[38]. The following ICD-10 diagnosis codes were used: ASD - F84.0, F84.1, F84.5, F84.8, F84.9; any psychiatric disorder – F00-F99. Reported diagnoses for the conditions used as exclusions were obtained from the DNPR, which holds all data on in- and out-patient diagnoses given at discharge from somatic wards in all hospitals and clinics since 1995[39]. **Supplementary Table 1** gives a full overview of relevant diagnosis codes. The MINERvA study was approved by the Regional Scientific Ethics Committee in Denmark and the Danish Data Protection Agency.

### DNA methylation profiling in MINERvA

Neonatal dried blood spot samples were retrieved from the Danish Neonatal Screening Biobank, within the Danish National Biobank, as part of the iPSYCH study. Neonatal DNA extractions and DNA methylation quantification was performed at the Statens Serum Institut (SSI, Copenhagen, Capital Region, Denmark), building on a previously described protocol[40]. Briefly, from each dried blood spot sample two disks of 3.2mm were used with the Extract-N-Amp Blood PCR kit (Sigma-Aldrich, St. Louis, United States of America) and eluted in 200μL buffer. 160μL of the isolated genomic DNA was converted with sodium bisulfite using the EZ-96 DNA Methylation Kit (Zymo Research, California, United States of America). DNA methylation was quantified across the genome using the Infinium HumanMethylation450k array (“450K array”) (Illumina, California, United States of America) and a modified protocol as previously described[39]. Fully methylated and unmethylated control samples were included on each plate throughout each stage of processing.

### MINERvA 450K array data pre-processing and quality control

Signal intensities for 1,316 neonatal blood samples, 14 fully methylated control samples and 14 fully unmethylated control samples were imported into the *R* programming environment using the *methylumIDAT*() function in the *methylumi* package[41]. Our stringent quality control (QC) pipeline included the following steps: 1) checking methylated and unmethylated signal intensities and excluding samples where either the median methylated or unmethylated intensity values were < 2500, 2) using the ten control probes to ensure the sodium bisulfite conversion was successful, excluding any samples with a median score < 80, 3) identifying the fully methylated and fully unmethylated control samples were in the correct location on each plate, 4) using the 65 SNP genotyping probes on the array to identify duplicate samples, 5) multidimensional scaling of data from probes on the X and Y chromosomes separately to confirm reported gender, 6) comparing genotype data for up to 65 SNP probes on the 450K array with SNP array data, 7) using the *pfilter*() function in *wateRmelon*[42] to exclude samples with more than 1% of probes characterized by a detection P-value > 0.05, in addition to probes characterized by > 1% of samples having a detection P-value > 0.05. In total, 1,263 samples (96.0%) passed all QC steps and were included in subsequent analyses. Normalization of the DNA methylation data was performed used the *dasen*() function in the *wateRmelon* package[42].

### SNP genotyping and derivation of ASD polygenic risk scores

DNA was extracted at SSI as above and whole genome amplified in triplicate using the REPLI-g kit (Qiagen, Hilten, Germany). The triplicates were pooled and then quantified using Quant-iT picogreen (Invitrogen, California, United States of America). Samples were genotyped at the Broad Institute (Boston, Massachusetts, United States of America) using the Infinium PsychChip v1.0 array (Illumina, San Diego, California, United States of America) using a standard protocol. Phasing and imputation was done using SHAPEIT[43] and IMPUTE2 with haplotypes from the 1000 Genomes Project, phase 3[44, 45] as described previously[37]. ASD polygenic risk scores (PRSs) were generated as a weighted sum of associated variants as previously described[46]. Briefly, results from the largest autism genome-wide association study (GWAS) available from a combined effort by the Psychiatric Genomics Consortium (PGC) and iPSYCH (Grove et al, https://www.biorxiv.org/content/early/2017/11/25/224774) was used to select genetic variants and provide weights. As the MINERvA cohort is a subset of the broader iPSYCH cohort we used GWAS results excluding MINERvA samples, so that there was no overlap between the training cohort and the test cohort. Ten different significance thresholds (p_T_) from 5×10^−8^ to 1 were used to select sets of genetic variants, which were LD clumped using *plink* with setting –*clump-p1 1 –clump-p2 1 –cump-r2 0.1 –clump-kb 500* to generate PRSs.

### Statistical analysis

All statistical analyses were performed using the R statistical environment version 3.2.2[47]. To validate the validity and robustness of our blood spot DNA methylation measures, we implemented two DNA methylation clock algorithms to derive estimates for both age in years[35] and gestational age in weeks[34] for each sample. In addition, for each sample, we computed a score for prenatal exposure to maternal smoking using DNA methylation data as previously described by Elliott et al[22]. To identify DNA methylation sites associated with ASD status in the MINERvA discovery dataset, a linear model was fitted for each DNA methylation site with DNA methylation as the dependent variable, case/control status as an independent variable and a set of possible confounders as covariates: sex, experimental array number, urbanicity level, birth month, birth year, gestational age, and cell composition variables estimated using the Houseman algorithm with a reference dataset for whole blood[48, 49]. Regional analysis was performed using the combP software[50]. Subsequent replication and meta-analysis was performed using summary statistics available from two U.S. based studies: the Study to Explore Early Development (SEED)[51] and the Simons Simplex Collection (SSC)[52]. A complete description of the SEED and SSC data can be found elsewhere [53]. Metaanalysis to combine the EWAS results from MINERvA, SEED, and SSC studies was performed for DNA methylation loci present in at least two of the three studies using Fisher’s method for combining P-values, focusing on DMPs where the same direction of effect was reported across all three studies. To identify DNA methylation sites associated with elevated autism polygenic risk burden, a linear model was used with DNA methylation as the dependent variable and ASD PRS, the number of nonmissing genotypes contributing to the PRS, the first five genetic principal components, sex, experimental array number, six cell composition variables, smoking score, gestational age, and birth weight included as independent variables as described above.

### DNA methylation quantitative trait loci (mQTL) and co-localization analyses

All DNA methylation sites located within 250kb of the three genome-wide significant genetic variants identified in the PGC-AUT GWAS [Grove et al, https://www.biorxiv.org/content/early/2017/11/25/224774] were identified and *cis* (defined as a 500kb window) mQTL analysis was performed using the 1,257 samples within MINERvA that had both DNA methylation and imputed genotype data. mQTL were identified using an additive linear model to test if the number of alleles (coded 0, 1, or 2) predicted DNA methylation at each site, including covariates for sex, and the first five principal components from the genotype data fitted using the MatrixEQTL package[54]. Co-localization analysis was performed for each DNA methylation site as previously described[55] using the R coloc package (http://cran.r-project.org/web/packages/coloc). From both the PGC-AUT GWAS data and our mQTL results we inputted the regression coefficients, their variances and SNP minor allele frequencies, and the prior probabilities were left as their default values. This methodology quantifies the support across the results of each GWAS for 5 hypotheses by calculating the posterior probabilities, denoted as *PPi* for hypothesis *Hi*.

> *H_0_: there exist no causal variants for either trait;*
>
> *H_1_: there exists a causal variant for one trait only, ASD;*
>
> *H_2_: there exists a causal variant for one trait only, DNA methylation;*
>
> *H_3_: there exist two distinct causal variants, one for each trait;*
>
> *H_4_: there exists a single causal variant common to both traits*.

## RESULTS

### Robust epigenetic signatures of chronological age and prenatal tobacco exposure validate DNA methylation data generated from neonatal blood spots

Following our stringent QC pipeline (see **Materials and Methods**) our final MINERvA DNA methylation dataset included 1,263 samples comprising 629 ASD cases and 634 controls. The characteristics of this sample are displayed in **Table 1**; of note, due to oversampling female cases, we had a near equal ratio of males and females (632:631). There were no significant differences between ASD cases and controls for maternal or paternal age, days to blood spot sampling, or birth weight (P > 0.05). There was a significantly higher rate of maternal smoking for the ASD cases (P = 0.003) and evidence of higher smoking quantity (P = 0.006). We used DNA methylation data to derive estimates of gestational age[34] and chronological age[35] for each sample. The mean predicted gestational age was 37.7 weeks (sd = 1.35 weeks; **Supplementary Figure 2**) compared to the actual mean of 39.6 weeks (sd = 1.77 weeks), with a strong positive correlation between estimated and actual gestational age (r = 0.602; **Figure 1A**). The mean predicted chronological age was 0.495 years (sd = 0.298; **Supplementary Figure 3**) and this was less strongly correlated with actual age (r = 0.139; **Figure 1B**), consistent with data from Knight et al[34]. Of note, “days to sampling” - i.e. the time between birth and blood draw - was not correlated with either predicted gestational age or chronological age, and controlling for this did not improve the strength of the correlation with gestational age (**Supplementary Figure 4**). We next tested robust markers of smoking exposure during pregnancy[23] and adulthood, using an established algorithm[22] to calculate a DNA methylation derived “smoking score” which we compared to reported *in utero* exposure. We identified a highly significant association between this smoking score and actual exposure, with offspring exposed to tobacco smoking in utero having higher smoking scores compared to offspring who were not exposed (P = 8.41 × 10^−95^) (**Figure 1C**)[22, 33]. Taken together these analyses show that neonatal blood spots can be used to generate reliable DNA methylation data that can robustly identify exposure-/trait-associated variation.

**Figure 1:**
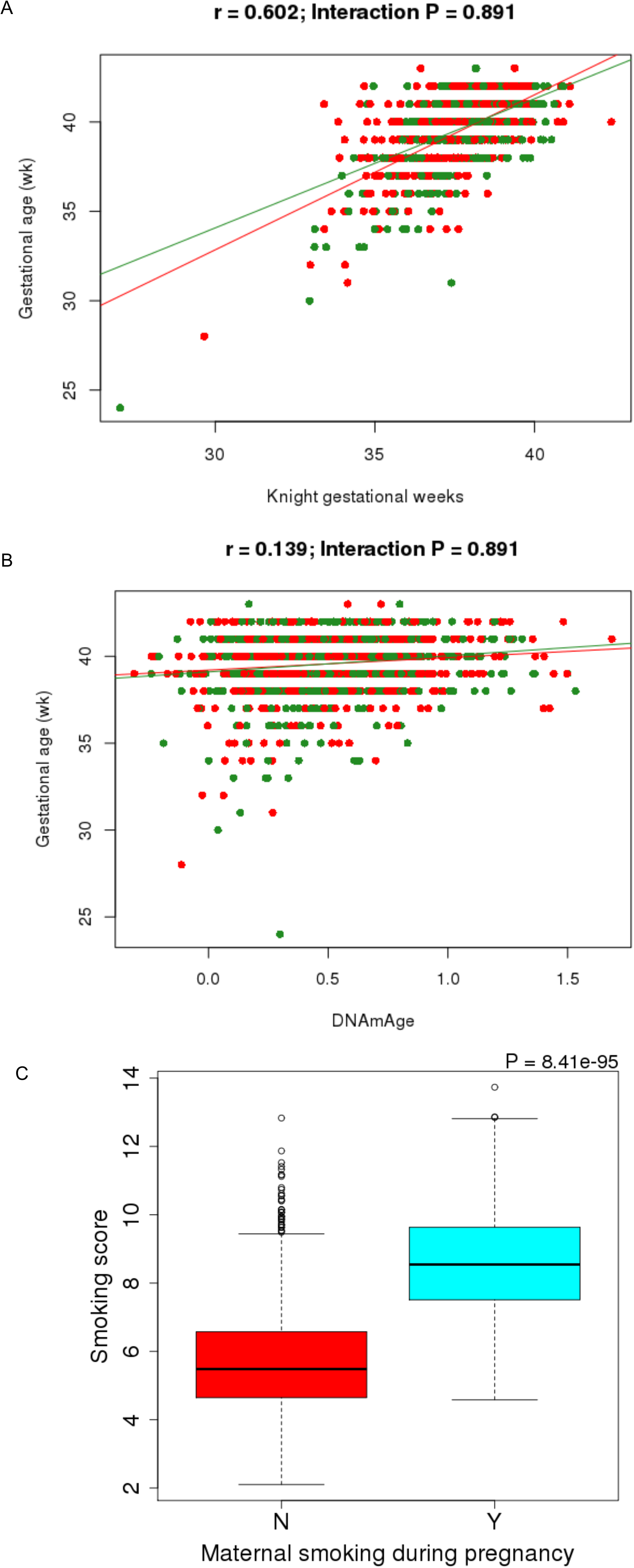
DNA methylation data from neonatal blood spots can be used to accurately predict age and maternal smoking status. A) Scatterplot of gestational age predicted from DNA methylation data (using an algorithm generated by Knight et al[34]) against actual gestational age. Autism cases are in red and controls are in green. B) Scatterplot of chronological age predicted from DNA methylation data (using the online Epigenetic Clock software[35]) against actual gestational age. Autism cases are in red and controls are in green. C) Boxplot of a smoking score derived from DNA methylation data[22] stratified by maternal smoking status during pregnancy.

**Table 1.** Characteristics of samples included in the MINERvA cohort. * = primary characteristics used to match cases and controls. ** = secondary characteristics used to match cases and controls as closely as possible.

| Characteristic | Unit/Category | ASD | Controls | P-value |
| --- | --- | --- | --- | --- |
| Sex * (%) | male | 52.0 | 49.8 | 0.933 |
| Birth year * (%) | 1998 | 31.8 | 30.6 | 0.991 |
|  | 1999 | 28.3 | 29 |  |
|  | 2000 | 7 | 6.78 |  |
|  | 2001 | 13.7 | 14.2 |  |
|  | 2002 | 19.2 | 19.4 |  |
| Gestational age ** (mean (SD)) | weeks | 39.6<br>(1.82) | 39.6<br>(1.72) | 0.96 |
| Urbanicity ** (%) | 1: Capital | 18.1 | 17 | 0.879 |
|  | 2: Suburb of the capital | 14.9 | 13.4 |  |
|  | 3: Municipalities having a town with more than 100,000 inhabitants | 8.27 | 8.68 |  |
|  | 4: Municipalities having a town with between 10,000 and 100,000 inhabitants | 27.3 | 29.2 |  |
|  | 5: Other municipalities in Denmark (largest town has less than 10,000 inhabitants) | 31.3 | 31.7 |  |
| Time to sampling (Mean (SD)) | days | 6.01<br>(3.15) | 6.15<br>(3.33) | 0.46 |
| Maternal age - Mean (SD) | years | 29.2<br>(4.94) | 29.7<br>(4.57) | 0.07 |
| Paternal age - Mean (SD) | years | 32.1<br>(6.04) | 31.9<br>(5.40) | 0.476 |
| Maternal smoking during pregnancy (%) | Smoke at any time | 29.0 | 21.2 | 0.00256 |
|  | Non-smoker | 71.0 | 78.8 |  |
| Maternal smoking amount during pregnancy (%) | 5 or less cigarettes per day | 6.46 | 7.25 | 0.00573 |
|  | 6-10 cigarettes per day | 11.2 | 7.59 |  |
|  | 11-20 cigarettes per day | 9.6 | 5.06 |  |
|  | 21 or more cigarettes per day | 1.05 | 1.18 |  |
| Birth weight - Mean (SD) | grams | 3512<br>(581) | 3541<br>(542) | 0.355 |

### Methylomic variation in perinatal blood is not significantly associated with childhood autism

Our initial analysis focused on identifying neonatal blood DNA methylation differences among MINERvA neonates who went on to develop a childhood diagnosis of ASD. Using a linear model to identify DNA methylation differences in ASD cases (N = 629) compared to controls (N = 634) we did not identify any differentially methylated positions (DMPs) passing a genome-wide significance threshold adjusted for multiple testing (P < 1×10^−7^). 20 ASD-associated DMPs were identified at a “discovery” threshold of P < 5×10^−5^ (**Supplementary Figures 5 & 6; Supplementary Table 2**); the most significant association was at cg12699865 which is located the 5’UTR of *RALY* where the mean level of DNA methylation was 0.647% lower (P = 7.63×10^−7^) in ASD cases (**Supplementary Figure 7**). Regional analysis combining the EWAS p-values for DNA methylation sites within a sliding window did not identify any significant ASD-associated differentially methylation regions. Given the higher prevalence of ASD diagnosis in males we tested for an interaction between autism status and sex but identified no significant associations (P < 1×10^−7^) and only seven DMPs at our discovery threshold of P< 5×10^−5^ (**Supplementary Table 3**).

We next meta-analyzed these findings with summary statistics from 450K array measurements for two U.S based studies of autism – the Study to Explore Early Development (SEED)[51] and Simons Simplex Collection (SSC)[52]. Although neither of these datasets was generated on blood samples collected immediately after birth, they enabled us to assess a combined sample size of 1,425 ASD cases and 1,492 controls (**Supplementary Table 4**). We first took the top ranked loci identified in each independent study and compared the directions of effect (i.e. difference between autism and controls); we did not find any excess of consistent associations (all sign test P > 0.05; **Supplementary Figure 8; Supplementary Table 5**). Second, we combined the p-values from the EWAS results of the three samples using Fisher’s Method (**Figure 2A & Supplementary Figure 9**). There were no sites where the combined P-value survived correction for multiple testing (P < 1×10^−7^), although 45 ASD-associated DMPs were identified at the discovery P-value threshold (P < 5 × 10^−5^) (**Supplementary Table 6**). The most significant DNA methylation site, based on a consistent direction of effect across all three studies, was cg03618918 (combined P = 3.85×10^−7^; pooled mean = 1.17%; **Figure 2B**), located ~10kb from *ITLN1*. In general, the estimated effects of ASD-associated DMPs (P < 5×10^−5^) was very small (**Supplementary Figure 10**), typically ~1% difference between ASD and controls. Taken together, these data suggest that ASD is not associated with robust methylomic signatures in blood obtained during early childhood.

**Figure 2:**
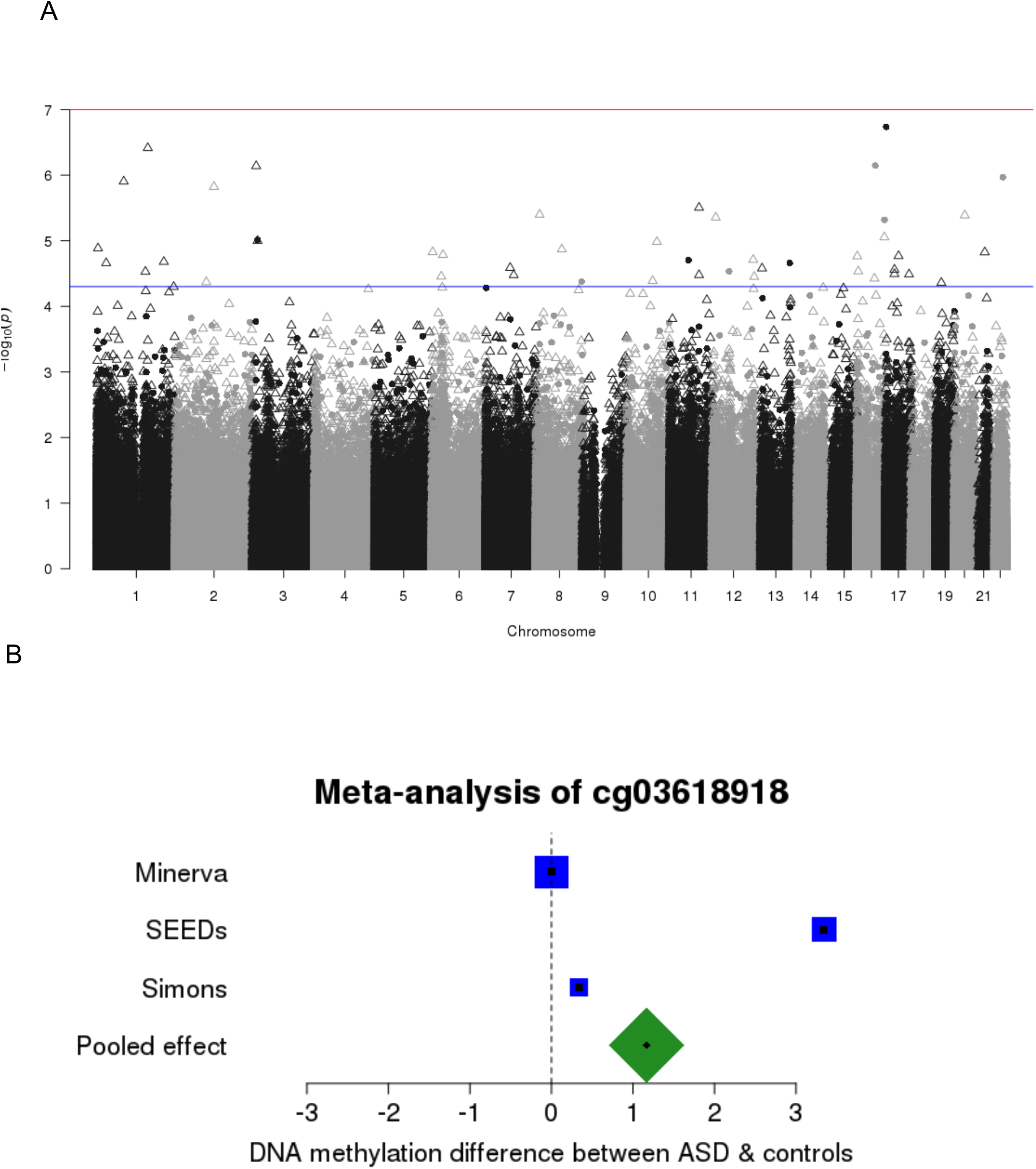
A cross-cohort meta-analysis finds little evidence of autism-associated methylomic variation in neonatal and childhood blood samples. A) Manhattan plot of P-values from the autism EWAS meta-analysis (total n = 2,917). P-values were calculated using Fisher’s method for combining P-values; solid circles indicate sites where the direction of effect was consistent across all contributing cohorts, empty triangles indicate where there were different directions of effect in at least two studies. The red horizontal line indicates experiment-wide significance (P < 1×10^−7^). B) Forest plot of cg03618918, the most significant DNA methylation sites associated with ASD in the meta-analysis. The effect is the mean difference in DNA methylation between autism cases and controls. The sizes of the boxes are proportional to the sample size of that cohort.

### Increased polygenic burden for autism is associated with methylomic variation in blood at birth

Like many complex diseases, individual genetic variants associated with autism explain only a tiny proportion of an individual’s risk[6, 56]. Polygenic risk scores (PRS), which essentially count the number of risk alleles across multiple associated loci, have been used successfully to capture the polygenic architecture of complex traits including autism[46]. PRS have been used to establish genetic correlations between traits[6] and there has been recent interest in using PRS as a quantitative variable to identify molecular biomarkers of high genetic burden[33, 57, 58]. PRS-associated epigenetic variation is potentially less affected by non-genetic risk factors for the disease itself which can confound case-control analyses, although pleiotropic effects of these genetic variants, which may themselves influence DNA methylation, cannot be excluded. We generated autism PRS for the iPSYCH-MINERvA sample using recent results from a meta-analysis of samples in PGC-AUT (Grove et al, https://www.biorxiv.org/content/early/2017/11/25/224774) excluding the subset of individuals included the MINERvA cohort (n = 45,162; 39.4% autism cases). Individual PRSs were calculated using a range of different GWAS P-value thresholds (p_T_ = 5×10^−8^, …, 1) to identify the optimal set of SNPs with the largest difference between ASD cases and controls in MINERvA. All scores based on P-values < 1 significantly predicted autism status (P < 0.05; **Supplementary Table 7, Supplementary Figure 11**), with a PRS based on p_T_ = 0.1 having the most significant difference (P = 9.49×10^−13^) between ASD cases and controls (**Figure 3A**). There was a strong positive correlation between scores based on SNPs selected at relatively relaxed significance thresholds (i.e. p_T_ > 0.001; **Supplementary Figure 12**), with weaker correlations between scores based on more limited (but more strongly associated) sets of variants, potentially reflecting the more dramatic effect a single SNP has on the PRS when the total number of SNPs is small. We next performed an EWAS of ASD PRS (**Supplementary Figure 13 and Supplementary Figure 14**), observing strong correlations (r > 0.5) between the results of analyses of scores based on p_T_ > 0.01 (**Supplementary Figure 15**). Examples of PRS-associated DMPs identified using the most predictive ASD PRS (p_T_ < 0.1) are shown in **Supplementary Figure 16**; in total, we identified two DMPs significantly associated (P < 1×10^−7^) with elevated polygenic burden (cg02771117: P = 3.14 × 10^−8^ and cg27411982: P = 8.38 × 10^−8^), with 49 DMPs associated at a more relaxed “discovery” P-value threshold (P < 5 × 10^−5^) (**Figure 3 and Supplementary Table 8**). Both cg02771117 and cg27411982 are located on chromosome 8, but are ~5kb apart and annotated to two different genes (FAM167A and RP1L1, respectively). Differential DNA methylation at these sites on chromosome 8 is identified in each of the eight most inclusive ASD PRS EWAS analyses (i.e. those using the most relaxed GWAS p-value threshold) (**Supplementary Figure 14**). Of note, both DMPs flank a significant genetic association signal identified in the latest ASD GWAS (**Supplementary Figure 17**). To establish whether the PRS-associated methylation signal in this region reflected direct effects of the GWAS signal itself, we iteratively added PRS variants within 100kb of these two sites as covariates in our EWAS in order of significance (see Materials and Methods). After the addition of the four most significant genetic variants, which were independently associated with cg02771117 (**Supplementary Figure 18**), the ASD PRS term was no longer significant (P = 0.0518; **Supplementary Table 9**). In contrast cg27411982 was still nominally significant even after the addition of 12 ASD associated SNPs, four of which however were independently associated and largely explained the association between the ASD PRS and DNA methylation (**Supplementary Figure 19; Supplementary Table 10**). These data suggest that the PRS-associated variation in DNA methylation at both cg02771117 and cg27411982 results from the combined effects of multiple genetic variants associated with ASD in this region. In order to demonstrate that the PRS EWAS results are not simply a consequence of the ASD cases within the full MINERvA sample, we repeated the analysis separately for cases and controls. P-values from this approach were strongly correlated with those for the analysis across all samples (**Supplementary Figure 20**), indicating that the methylomic consequences of high genetic burden are consistent across both groups.

**Figure 3:**
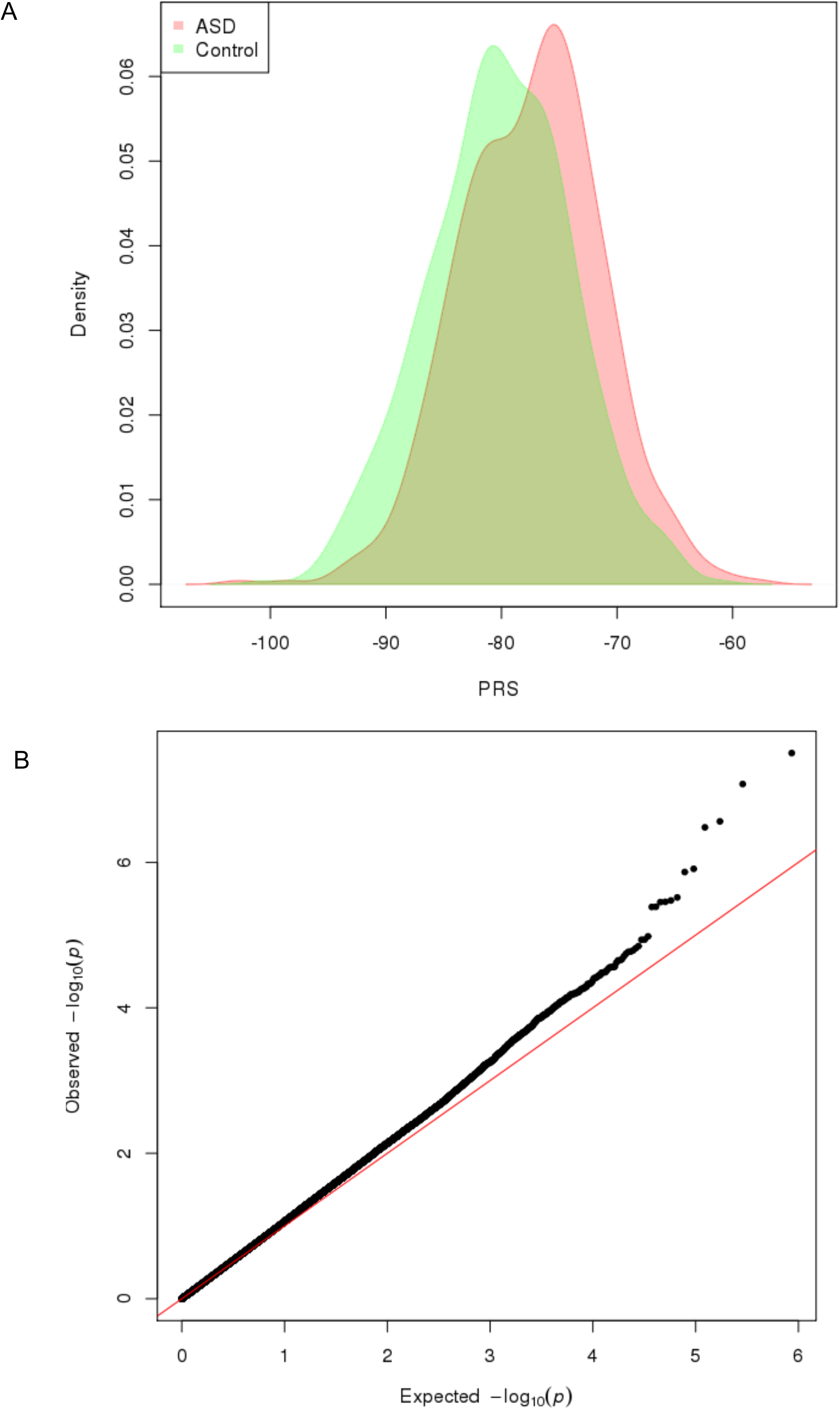
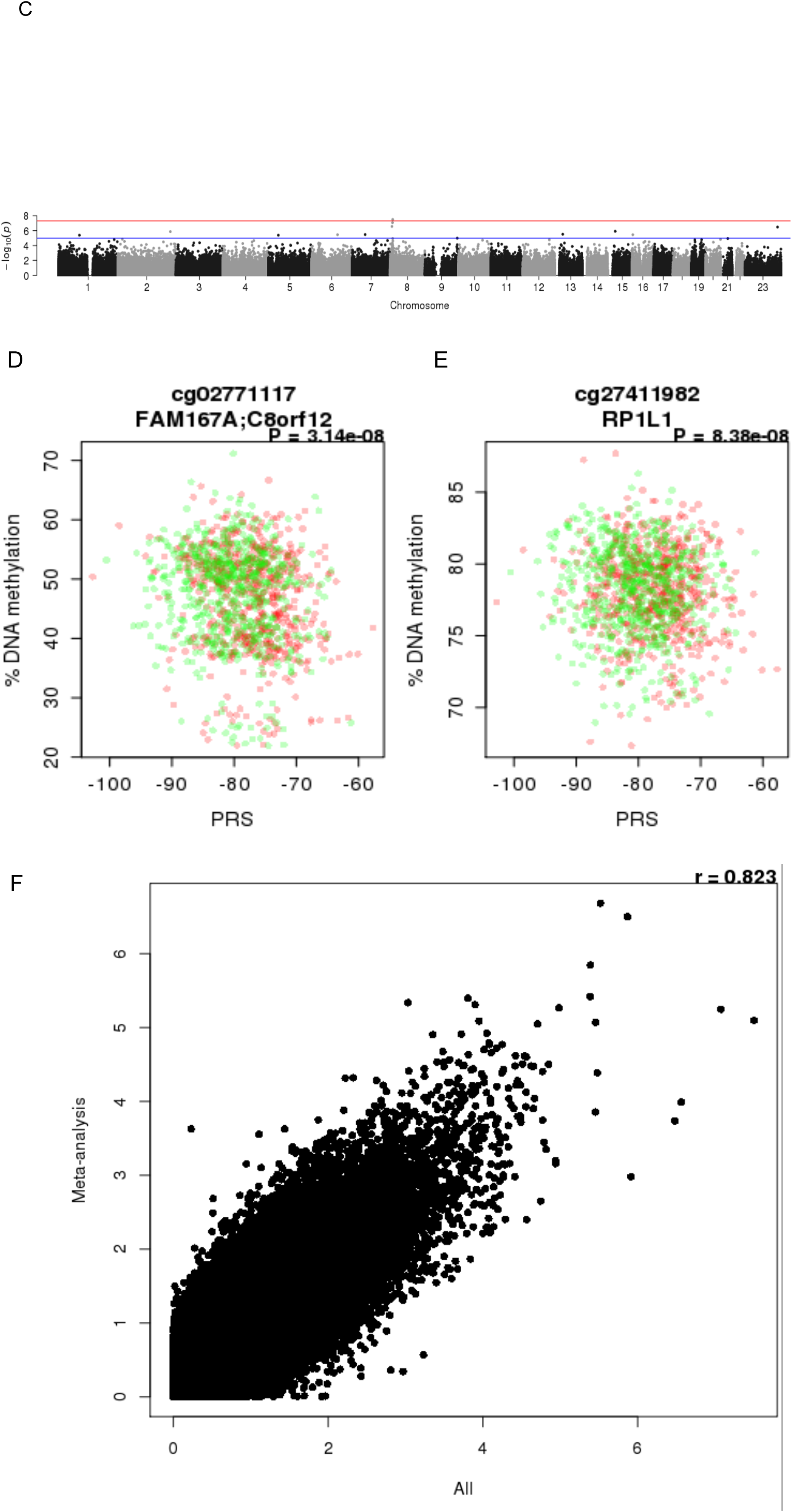
Polygenic burden for autism is associated with significant variation in DNA methylation at birth. A) Density plot of polygenic risk score (PRS) (p_T_ = 0.01) split by ASD case control status. B) Q-Q plots of the ASD PRS (p_T_ = 0.01) EWAS analysis in neonatal blood DNA. C) Manhattan plot of the ASD PRS (p_T_ = 0.01) EWAS analysis in neonatal blood DNA. The red horizontal line indicates experiment-wide significance (P < 1×10^−7^); blue horizontal line indicates a ‘discovery’ significance threshold (P < 5×10^−5^). Scatterplots of genome-wide significant CpG sites where DNA methylation (y-axis) at D) cg02771117 and E) cg27411982 is correlated with ASD PRS (x-axis). Red points indicate ASD cases, green points indicate controls. F) Scatterplots of –log10 P-value from the EWAS of ASD PRS comparing the results from an analysis performed in all individuals (x-axis) against the results from an analysis performed separately for cases and controls and then combined with a meta-analysis (y-axis).

### Alignment of DNA methylation quantitative trait loci and ASD genetic signals

None of the GWAS-AUT identified variants tag known nonsynonymous mutations; consistent with other complex phenotypes it is likely that disease-associated variants instead influence the regulation of gene expression[13, 59]. Building on our previous work showing how DNA methylation quantitative trait loci (mQTLs) can be used to refine GWAS loci through the identification of discrete sites of variable methylation associated with disease risk variants[29, 33] we used the matched MINERvA DNA methylation and genetic data (see **Materials and Methods**) to identify mQTL located in the vicinity of ASD-associated GWAS variants (**Figure 4, Supplementary Figure 21**). Simply aligning mQTL data with GWAS results is not sufficient to infer that there is relationship between ASD and DNA methylation in these regions; instead it may reflect two distinct causal variants - one associated with ASD and the other with DNA methylation - in strong linkage disequilibrium. To establish whether there was evidence of a single causal variant influencing both DNA methylation and ASD in the regions nominated by the GWAS we performed a Bayesian colocalization analysis[55]. Briefly, this approach compares the pattern of association results from two independent GWAS (i.e. of ASD and DNA methylation) to see if associations colocalize to the same causal variant. We considered mQTL data for 457 unique Illumina 450K probes located within 250kb of three independent autosomal ASD GWAS variants. The posterior probabilities involving 91 DNA methylation sites are supportive of a co-localized association signal for both ASD and DNA methylation (PP_3_+PP_4_ > 0.99; **Supplementary Table 11**). Four of these sites located on chromosome 20 had a higher posterior probability for both ASD and DNA methylation being associated with the same causal variant compared to them being associated with different causal variants (PP_4_/PP_3_ > 1; **Supplementary Figure 22; Supplementary Figure 23**) and the genes annotated to these sites (KIZ, XRN2, and NKX2-4) represent putative candidates for a potential functional role in ASD.

**Figure 4:**
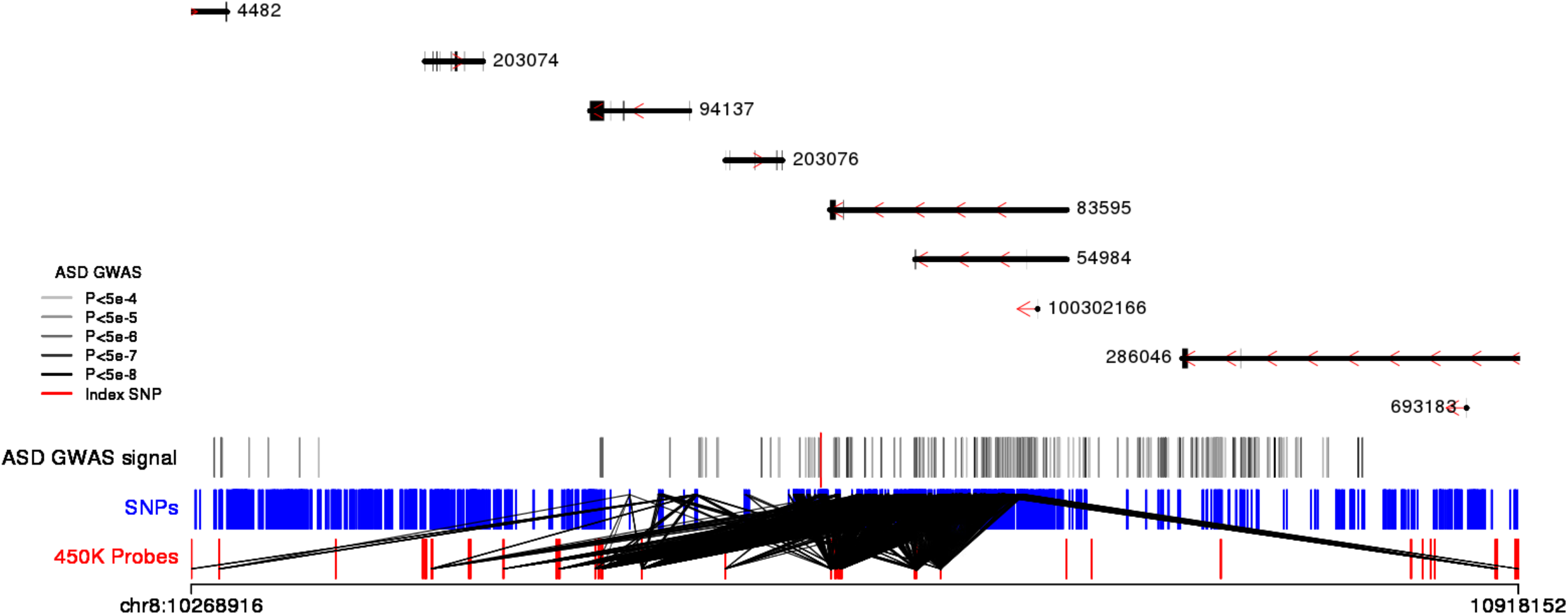
DNA methylation quantitative trait loci (mQTL) mapping can localize putative causal loci associated with ASD. Presented here is a genomic region (chr8:10268916-10918152) identified in a recent GWAS analysis of ASD [Grove et al, https://www.biorxiv.org/content/early/2017/11/25/224774]. At the top of the figure is a schematic detailing the genes located in this region which are identified by their Entrez ID number. All genetic variants identified in the ASD GWAS (P < 1×10^−4^) are represented by vertical solid lines where the color reflects the strength of the association ranging from gray (less significant P-values) to black (more significant P-values). A red vertical line indicates the most significant genetic variant in this region. All DNA methylation sites tested for neonatal blood mQTL in the MINERvA dataset are indicated by red vertical lines and genetic variants by blue vertical lines. Significant neonatal blood mQTLs (P < 1×10^−13^) are indicated by black diagonal lines between the respective genetic variant and DNA methylation site. Additional examples of mQTLs in genomic regions showing genome-wide significant association with ASD are given in **Supplementary Figure 21**.

## DISCUSSION

In this study, we quantified neonatal methylomic variation in 1,263 infants selected from the iPSYCH cohort[37] including samples from individuals who went on to develop ASD and carefully-matched control samples. It represents the first attempt to integrate analyses of both genetic and epigenetic variation in ASD, demonstrating the utility of using a polygenic risk score to identify molecular variation associated with disease, and of using DNA methylation quantitative trait loci to refine the functional and regulatory variation associated with ASD risk variants. While ASD itself was not associated with significant differences in neonatal DNA methylation, at a genome-wide significance threshold, increased polygenic burden for autism was found to be associated with methylomic variation at specific loci in blood at birth. Our analysis of ASD PRS and DNA methylation supplements an increasing body of literature investigating the effects of high genetic burden for other complex traits on molecular variation[33, 57, 58]. We find that two CpGs located on chromosome 8 are associated with genetic risk for ASD, and are proximal to a robust GWAS signal for ASD. Furthermore, multiple associated SNPs on chromosome 8 have a polygenic effect on DNA methylation at these two CpG sites, demonstrating how a complex genetic architecture can converge on a common molecular consequence.

This study has several advantages over previous analyses of DNA methylation in ASD. We assessed a relatively large set of samples that is balanced with regard to both disease status and numbers of males and females. This contrasts with previous studies that have been undertaken on much smaller numbers of samples and focused primarily on ASD in males. Our control samples were stringently matched to cases on the basis of a number of criteria (see **Methods**) to minimize the effects of confounding variables that often lead to false positives in molecular epidemiology. Furthermore, our use of neonatal DNA samples - collected before diagnosis and the manifestation of any ASD symptoms - means that we are uniquely positioned to identify epigenetic variation associated with later disease or elevated polygenic burden for later ASD, avoiding the confounding exposures often associated with disease (for example, medication, stress, and reverse causation)[60]. Finally, our study profiled whole blood from neonatal infants rather than cord blood; this minimizes confounding by maternal blood DNA and means our data can be more easily compared to blood datasets derived from later in life.

We find little evidence to support an association between DNA methylation at birth and ASD, confirming this finding in a meta-analysis of three studies with a total sample of 2,917. Power calculations show that we have >90% power in our meta-analysis to identify a ASD-associated difference of 0.3% and a difference of 0.7% in the MINERvA cohort alone. While this suggests the lack of association was not due to sample size, we cannot fully conclude that DNA methylation is not associated with the onset of ASD. First, our analyses were constrained by the technical limitations of the Illumina 450K array which only assays ~ 3% of CpG sites in the genome. Second, this work necessitated the use of a peripheral tissue that may provide limited information about variation in the presumed tissue of interest, i.e. the brain[61]. This is a salient point for understanding the role DNA methylation plays in the disease process; however biomarkers, by definition need to be measured in an accessible tissue and therefore justify the use of blood from neonates in this study. Third, given the chronology of sample collection prior to ASD diagnosis, it is plausible that we were looking too early on in the disease process. Another limitation of our study is the possibility of diagnostic misclassification, however validation of select diagnoses (e.g., schizophrenia, single-episode depression, dementia, and childhood autism) has been performed with good results[38, 62].

In contrast, we find that polygenic burden for ASD is robustly associated with DNA methylation at two CpG sites on chromosome 8, with 49 DMPs associated with ASD polygenic burden at a more relaxed “discovery” P-value threshold. Of note, both sites flank a significant genetic association signal identified in the latest ASD GWAS and our data suggest that the PRS-associated variation at these sites results from the combined effects of multiple genetic variants associated with ASD in this region. Finally, we have used mQTL analyses to annotate this extended genomic region nominated by GWAS analyses of ASD, using co-localization analyses to highlight potential regulatory variation causally involved in disease. Of interest, we found evidence that several SNPs on chromosome 20 were causally-associated with both ASD and DNA methylation and the genes annotated to these sites (KIZ, XRN2, and NKX2-4) represent putative candidates for a potential functional role in ASD.

## CONCLUSIONS

In summary, our data provide evidence for differences in DNA methylation at birth associated with an elevated polygenic burden for ASD. Our study represents the first analysis of epigenetic variation at birth associated with autism and highlights the utility of polygenic risk scores for identifying molecular pathways associated with etiological variation.

## DECLARATIONS

### Ethics approval

The MINERvA study has been approved by the Regional Scientific Ethics Committee in Denmark, the Danish Data Protection Agency and the NBS-Biobank Steering Committee.

### Data availability

Given the nature of the MINERvA cohort, access to data can only be provided through secured systems which comply with the current Danish and EU data standards. To comply with the study’s ethical approval, access to the raw data is only available to qualified researchers upon request. All summary statistics and analysis scripts are available directly from the authors (please contact Jonas Grauholm at).

